# Proteomic responses under differing pH and pCO_2_ levels in the diatom *Thalassiosira pseudonana* are consistent with a hybrid carbon concentrating mechanism

**DOI:** 10.1101/2025.06.11.659125

**Authors:** Anthony R. Himes, Adam B. Kustka

**Affiliations:** Department of Earth and Environmental Sciences, Rutgers University—Newark, 101 Warren Street, Newark, NJ 07102, USA

**Keywords:** acid-base regulation, alternative oxidase, CCM, carbon uptake, carbonate chemistry, carbonic anhydrase, malate dehydrogenase, malate/oxaloacetate shuttle, phosphoenolpyruvate carboxylase, proteome

## Abstract

The effects of ongoing anthropogenic climate change are not well known in marine diatoms, a key group of primary producers. In particular, detailed characterizations of their carbon concentrating mechanisms (CCM) are lacking, which limits the understanding of how changing ocean carbonate chemistry will impact global primary production. While the model diatom *Thalassiosira pseudonana* has been widely studied, contrasting results have prevented the clear elucidation of its CCM. A quantitative proteomic analysis was therefore performed across three experimental treatments (low pCO_2_/high pH, high pCO_2_/low pH, low pCO_2_/low pH) to discern the specific roles of proteins that can be involved in CCMs as well as other cellular processes (e.g. pH and redox regulation). This analysis revealed a hybrid CCM consisting of both biophysical and biochemical steps that facilitate increased CO_2_ diffusion into the cell, the formation and transport of an organic carbon intermediate into the chloroplast, the subsequent decarboxylation of this intermediate, and the facilitated diffusion of inorganic carbon from the stroma into the pyrenoid-penetrating thylakoid. No evidence supporting alternative roles for the identified CCM proteins was found. As several aspects of this CCM require further investigation, common challenges inherent to CCM research are discussed and strategies to overcome them are suggested.

**Highlight:** Experimental manipulation of carbonate chemistry allowed for the separation of the carbon concentrating mechanism in *T. pseudonana* from other processes that rely on the same proteins, revealing a hybrid CCM.

## Introduction

In the face of ongoing anthropogenic climate change, the need to further our understanding of how various taxonomic groups will be impacted is of ever-increasing importance. Marine phytoplankton are one key group, as they account for about half of all global primary production (Behrenfeld *et al*., 2001; Westberry *et al*., 2023). In particular, diatoms account for the largest proportion of marine planktonic primary production, with estimates ranging from 20 to 50% (Neslon *et al*., 1995; Field *et al*., 1998; Armbrust, 2009; Rousseaux and Gregg, 2014; Tréguer *et al*., 2018). As anthropogenic carbon dioxide (CO_2_) emissions continue to drive ocean acidification, the concomitant increase in oceanic partial pressure of CO_2_ (pCO_2_) and decrease in pH will directly affect dissolved inorganic carbon (DIC) speciation, and therefore, have direct effects on all carbon fixing organisms (IPCC, 2023). However, the nature of these impacts are still unclear within diatoms as their mechanisms for DIC uptake and its subsequent transport through the cell to ribulose-1,5-bisphosphate carboxylase/oxygenase (RuBisCO), known collectively as carbon concentrating mechanisms (CCM), are not well understood (Granum *et al*., 2005; Trimborn *et al*., 2008; Reinfelder, 2011; Kustka *et al*., 2014; Zhang *et al*., 2014; Clement *et al*., 2016; Wu *et al*., 2021; Nam *et al*., 2024; Shimakawa *et al*., 2024).

There are two classes of CCMs broadly termed biophysical and biochemical based on whether a given mechanism relies solely on bicarbonate (HCO_3_^-^) pumping and CO_2_ diffusion or includes the formation, transport, and decarboxylation of an acid-stable four carbon intermediate, respectively (see review by Reinfelder, 2011). Both concentrate CO_2_ near RuBisCO, which in diatoms is localized to the pyrenoid to minimize photorespiratory losses (Edwards *et al*., 2004; Mackinder *et al*., 2016; Nam *et al*., 2024; Shimakawa *et al*., 2024). Biophysical CCMs in marine phytoplankton depend on the active transport of extracellular HCO_3_^-^ through multiple subcellular compartments alongside various forms of carbonic anhydrase (CA) that help to prevent the diffusive loss of CO_2_ (Tachibana *et al*., 2011; Samukawa *et al*., 2014; Tanaka *et al*., 2014; Hennon *et al*., 2015; Nawaly *et al*., 2023b; Shimakawa *et al*., 2023).

For biochemical CCMs, much understanding stems from vascular land plants, where C_4_ plants are known to store inorganic carbon as four-carbon compounds before transport near to RuBisCO, and subsequent decarboxylation to release CO_2_. One commonly observed C_4_ plant CCM begins with phosphoenolpyruvate carboxylase (PEPC) converting cytosolic phosphoenolpyruvate (PEP) and CA-derived HCO_3_^-^ into oxaloacetate (OAA). This OAA is then reduced to malate by malate dehydrogenase (MDH) before transport into the chloroplast where NADP^+^-dependent malic enzyme (ME) decarboxylates malate to pyruvate, NADPH + H^+^, and CO_2_. As RuBisCO is found within the chloroplast stroma of C_4_ plants, the decarboxylation of malate concludes this form of CCM. Evidence from several studies also supports the existence of similar biochemical CCMs within some diatoms (Reinfelder *et al*., 2004; Kustka *et al*., 2014; Wu *et al*., 2021; Yu *et al*., 2022; Zhen *et al*., 2025); however, in all cases, key differences have been observed between C_4_ plant and diatom CCMs. One major difference recently noted in two model diatom species, *Thalassiosira pseudonana* and *Phaeodactylum tricornutum*, is the presence of a specialized protein coat surrounding the pyrenoid that may act as a diffusion barrier to CO_2_, thereby requiring further CA-mediated steps to ultimately concentrate CO_2_ near RuBisCO (Nawaly *et al*., 2023b; Nam *et al*., 2024; Shimakawa *et al*., 2024).

Multiple studies have attempted to characterize the CCM of *T. pseudonana*, but have yielded inconsistent results with contrasting interpretations. (Reinfelder *et al*., 2004; Roberts *et al*., 2007; Granum *et al*., 2009; Trimborn *et al*., 2009; Kustka *et al*., 2014; Tanaka *et al*., 2014; Clement *et al*., 2016; Hopkinson *et al*., 2016). First, seemingly contradictory results on pCO_2_-dependent changes in HCO_3_^-^ uptake have been reported for *T. pseudonana* (Trimborn *et al*., 2009; Clement *et al*., 2016), although there were large differences between the “high” pCO_2_ levels used in these studies. Second, relative transcript abundances for the two known PEPC genes (PEPC1 and PEPC2) have shown inconsistent sensitivities to steady-state pCO_2_ conditions (Granum *et al*., 2005; Roberts *et al*., 2007; Tanaka *et al*., 2014) compared to abrupt shifts from higher to lower pCO_2_ levels (Granum *et al*., 2009; Kustka *et al*., 2014) leading to debate over the classification of the CCM of *T. pseudonana* as biophysical or biochemical. Third, while PEPC1 and PEPC2 protein abundances did not increase in line with their respective transcript levels immediately following a shift from high to low pCO_2_ (Granum *et al*., 2009), they did increase in response to steady-state low pCO_2_ conditions (Kustka *et al*., 2014), which highlights that gene expression levels are not always representative of cellular physiology. Lastly, in vitro PEPC to RuBisCO activity ratios have been used to argue in favor of a biophysical CCM (Trimborn *et al*., 2009; Clement *et al*., 2016), although it remains unclear how representative these methodologies are of true in vivo PEPC activity in *T. pseudonana*.

One difficulty of elucidating a CCM is the accurate localization of its protein components. For example, early predicted localizations of PEPC2 placed it within the mitochondrial matrix of diatoms (Kroth *et al*., 2008). Recent findings in other eukaryotic phototrophs, however, place it on the surface of mitochondria (O’Leary *et al*., 2011; Park *et al*., 2012; O’Leary and Plaxton, 2020), which is the same localization supported by current predictive tools for *T. pseudonana* (Ødum *et al*., 2024). Accurate protein localizations are but one of the prerequisites to adequately understand protein function.

Several metabolic pathways are known that, if upregulated in response to changing pCO_2_ and/or pH conditions, could be misinterpreted as evidence of a C_4_ CCM. For example, upregulation of PEPC under low pCO_2_/high pH conditions might be indicative of the intracellular pH homeostasis pathways described in Sakano (1998) where PEPC activity serves to acidify the cell. This acidification effect is then spread across subcellular compartments by the malate/OAA shuttle, which is the term given to the reduction of OAA to malate by MDH, the transport of malate across intracellular membranes, and its subsequent oxidation back to OAA by MDH (reviewed in Selinski and Scheibe, 2018). The malate/OAA shuttle is also known to be utilized to maintain cellular redox poise, facilitate cyclic electron transport, and dissipate excess light energy (Shikanai, 2007; Yoshida *et al*., 2007; van Dongen *et al*., 2011; Yu *et al*., 2014; Nawrocki *et al*., 2015; Dietz *et al*., 2016; Murik *et al*., 2018; Selinski and Scheibe, 2018; Chadee *et al*., 2021; Zhou *et al*., 2021; Hippmann *et al*., 2022; Degen *et al*., 2024). However, none of these alternative mechanisms have been evaluated simultaneously with the CCM of *T. pseudonana*.

Given the debate surrounding the identity of the CCM in *T. pseudonana*, this study employed a quantitative proteomic approach across three experimental conditions to assess each step constituting the CCM of this species. The three conditions evaluated were 1) low pCO_2_/high pH, 2) high pCO_2_/low pH, and 3) low pCO_2_/low pH. While *T. pseudonana* is unlikely to experience the third condition in a natural setting, as it was achieved by decreasing the total DIC pool, the inclusion of this treatment will help to disentangle any potential biochemical CCM from the aforementioned alternative mechanisms involving PEPC and the malate/OAA shuttle driven by changes in pH alone, irrespective of pCO_2_. In this way, this study aims to clarify and deepen the understanding of both the CCM of *T. pseudonana* as well as various other cellular pathways that may be modulated in response to increases in pCO_2_ and/or decreases in pH, thus providing insight into how diatoms as a whole will respond to ongoing ocean acidification.

## Materials and methods

### Stock culture conditions

*Thalassiosira pseudonana* (strain CCMP 1335) was maintained in a culture incubator (VWR, Radnor, PA, USA; Model No. 2015) set to 18°C with 24 h high-intensity light (250 μmol photons m^-2^ s^-1^). Precision incubator thermometers (VWR) were maintained alongside cultures and checked daily to ensure temperatures remained consistent. Stock cultures were grown in metal-replete Aquil* media following Sunda *et al*., 2005, with the Fe concentration lowered to 600 nM and the treatment with Chelex-100 omitted. Synthetic seawater was combined with macronutrient, trace metal, and vitamin stock solutions (Sunda *et al*., 2005) in clean sterile polycarbonate bottles (see supplemental *Cleaning and sterilization*) within a class 100 laminar flow hood (Mystaire, Creedmoor, NC, USA). Assembled media was vacuum filter sterilized using autoclaved 0.2 μm Supor PES membrane filter discs (Pall Corporation, Port Washington, NY, USA). *T. pseudonana* was then inoculated into sterile media contained within clean sterile 30 mL polycarbonate tubes. Semi-continuous batch cultures were maintained in this way for the duration of the study so that cells in logarithmic growth phase were readily available for experimentation. Growth of stock cultures was monitored daily by measuring relative chlorophyll a fluorescence using a Turner Designs 10-AU fluorometer (Turner Designs, San Jose, CA, USA). Growth rates were calculated from linear regressions of the natural logarithm of blank corrected relative chlorophyll a fluorescence over time in days.

### Experimental culture media

*T. pseudonana* was exposed to three experimental treatments: a low pCO_2_/high pH condition (pCO_2_ = 202 μatm, pH = 8.48), a high pCO_2_/low pH condition (pCO_2_ = 1702 μatm, pH = 7.61), and a low pCO_2_/low pH condition (pCO_2_ = 202 μatm, pH = 7.61). Hereafter these treatments will be referred to as LCHpH, HCLpH, and LCLpH, respectively. The letters H and L denoted high or low, pCO_2_ is shortened to the letter C, and pH is left unabbreviated. For each of the three treatments, metal-replete Aquil* was made following the same procedures outlined above, but with two key differences.

First, the concentration of NaHCO_3_ in the LCLpH treatment was lowered from 2.38 mM to 0.283 mM so that at pH 7.61 the pCO_2_ of this media would be equal to that of the LCHpH media. CO2SYS (Lewis and Wallace, 1998; see supplemental *CO2SYS constants*) was used to determine this lower DIC concentration while producing three treatments from two unique pH and pCO_2_ levels that would serve to distinguish the impact of pH alone from the CCM of *T. pseudonana*. To prevent carbon limitation in LCLpH cultures while also limiting the potential for fluctuations in carbonate chemistry driven by high culture biomass (Shi *et al*., 2009; Crawfurd *et al*., 2011), all experimental cultures were sampled at relatively low biomass (∼70,000 cells per mL^-1^).

The second key difference in the experimental media was that the total concentration of each trace metal varied among treatments. This was done so that free metal ion concentrations would be equivalent across treatments despite their differences in carbonate chemistry (Sunda *et al*., 2005). The total metal concentrations for each treatment were calculated using Visual MINTEQ ver. 4.0.

Once media was fully assembled and filter sterilized, a small aliquot was removed to measure pH using a freshly calibrated Oakton double-junction all-in-one pH/ATC electrode (Cole-Parmer 35811-72) connected to an Oakton pH 110 series pH meter (Cole-Parmer 35615-20). Dilute NaOH and HCl solutions, made in ultrapure water and 0.2 μm filter sterilized, were used to adjust culture media to the desired pH for each condition.

### Experimental apparatus

Experiments were carried out in modified 500 mL polycarbonate bottles. In brief (see supplemental *Experimental apparatus continued* for full details), three modifications were made to enable real-time pH monitoring and control: 1) an acid line was plumbed in through the neck of the bottle to deliver 0.006 M HCl, prepared in treatment-specific synthetic ocean water, to counteracted pH increases driven by photosynthetic activity. 2) An airline was run to the bottom of the bottle through which a mixture of ultra zero grade air and CO_2_ (Airgas, Radnor, PA, USA) was bubbled to mix cultures during acid additions. High accuracy mixer rotameters (Matheson Tri-gas Inc., Montgomeryville, PA, USA) were used to ensure each experimental culture was bubbled with air matching its target pCO_2_. 3) A Thermo Orion double junction pH electrode (Thermo Scientific 9156DJWP) was inserted through the cap of the bottle and was used to measure pH in real time throughout the course of each experimental trial. In each of three identical experimental systems, Alpha pH 200 series pH controllers (Eutech Instruments ECPHCTP0200) were used in conjunction with gas solenoid valves (ASCO U8256B045V) for air delivery and small peristaltic pumps (Masterflex C/L 77120-20) for acid delivery to maintain pH setpoints (similar to the setup used in Kustka *et al*., 2014). These controllers were programmed to maintain pH within 0.02 pH units of the target level, which matched the reported precision for the pH probes. During testing, pH probe drift over the course of one week was negligible; therefore, pH probes were freshly calibrated at the start of each 4-day experimental trial. Culture pH was recorded twice daily for the duration of each trial.

A second air delivery system was used to refresh the headspace of each culture bottle with the appropriate pCO_2_ gas mixture to minimize laboratory air intrusion as bottles could not be kept sealed while actively regulating pH. Solenoid valves were programmed to open every 15 min for a 2-min duration using outdoor plugs (Wyze Labs, Inc., Seattle, Washington, USA). During testing, this additional air turnover was found to markedly improve pH control.

### Experimental trials

At the start of each experimental trial, 450 mL of sterile media was inoculated with mid-logarithmic growth phase *T. pseudonana*. Three cultures, one of each treatment group, were always run in parallel within the same incubator that was used to maintain stock cultures. Trials (n = 8) lasted for approximately 4 d and growth was monitored daily by measuring cell counts and biovolumes with a Multisizer 3 Coulter counter (Beckman Coulter Life Sciences, Indianapolis, IN, USA). Specific growth rates (μ) were then calculated from the linear regression of the natural logarithm of cell density (cell mL^-1^) over time in days (Table S2). Experimental cultures were inoculated such that a minimum of 10 cell divisions occurred prior to sampling. Once cultures reached approximately 70,000 cells mL^-1^, cells were collected via vacuum filtration onto 3 μm polycarbonate filter discs (MilliporeSigma). Cells were then transferred into microcentrifuge tubes and flash frozen in liquid nitrogen before storage at -80°C for later analysis.

### Protein extraction and quantification for proteomics

Cell pellets were thawed on ice and bead beaten using lysing matrix B (MP Biomedicals, Irvine, CA) in buffer containing 50 mM HEPES pH 8.5, 1% SDS, and protease inhibitor cocktail (MilliporeSigma, P2714). A FastPrep-24 5G bead beating lysis system (MP Biomedicals) was used with a speed of 4 m s^-1^ for two 30 s cycles. Cellular debris was then removed by centrifugation at 16,000 g for 5 min. As the proteomic sample processing method used in this study is sensitive to interference by genomic DNA released during cell lysis, cell extracts were next incubated for 25 min at 4°C with end-over-end rotation in the presence of 500 units mL^-1^ Benzonase nuclease (MilliporeSigma, E1014). A small volume of each sample was then removed for protein quantification before 10 mM dithiothreitol (DTT) was added to the remaining lysate. Samples were frozen at -80°C for later analysis.

Total protein concentrations were determined in duplicate for each sample using the Micro BCA protein assay kit (Thermo Scientific, 23235) following the manufacturers specifications. Diatom stoichiometric ratios of cellular carbon and nitrogen to protein content (Lourenço *et al*., 1998; Lourenço *et al*., 2004) were used along with measured cell densities and cell biovolumes to estimate the expected protein yield from each sample. Quantified total protein concentrations were compared to these estimates and reported as protein extraction efficiencies in Table S2. The four samples within each treatment with the highest extraction efficiencies were selected for proteomic sequencing. Proteomic sample preparation following the SP3 method (Hughes *et al*., 2014) and LC-MS/MS sequencing were completed by staff at the Biological Mass Spectrometry Facility of Robert Wood Johnson Medical School and Rutgers, The State University of New Jersey (see supplemental *Proteomic sample processing* for details).

### Proteomic data analysis and statistics

LC-MS/MS output files were produced in Thermo .raw format and were processed using DIA-NN ver. 2.0.2 (Demichev *et al*., 2020; Kistner *et al*., 2023, Preprint). In silico spectral library preparation began with curating available protein sequences from UniProt for *T. pseudonana*, which yielded 11,661 unique protein entries (see supplemental *In silico spectral library preparation* for details). These sequences were then used in conjunction with a universal contaminants library (Frankenfield *et al*., 2022) to generate the in silico spectral library needed for protein identification and quantification. For the sample analysis run, the high precision Quant UMS quantification strategy was used with match between runs (MBR) enabled, protein inference enabled, and the precursor false discovery rate (FDR) set to 1%. The main output report was then imported into R ver. 4.4.1 (R Core Team, 2024) for strict quality control filtering before robust ridge regression model fitting using the MSqRob2 Bioconductor R package ver. 1.12.0 (Goeminne *et al*., 2016; Goeminne *et al*., 2020; Sticker *et al*., 2020). The full list of DIA-NN settings and filters are provided in the supplemental section *Data analysis with DIA-NN*. All pairwise comparisons between the LCHpH and HCLpH treatments and the LCHpH and LCLpH treatments were evaluated using the Benjamini-Hochberg method to control FDR (Tables S5 and S6). Differential protein abundances greater than 1.30-fold with an adjusted P < 0.05 were considered significant.

### Predicted localizations of key proteins

Sequence analysis to explore the specific mitochondrial association of PEPC2 was completed using DeepLoc 2.1 set to high quality (slow) and long output (Ødum *et al*., 2024). ASAFind 2.0 (Almagro Armenteros *et al*., 2019; Gruber *et al*., 2025) was used to reassess the predicted chloroplast localization of the one form of PYC detected in this study as it was specifically developed for use in diatoms.

## Results

### Culture conditions and growth

Extensive testing was conducted to achieve suitable pH control of experimental cultures (n=13-15 per treatment, data not shown). The first iteration of the experimental apparatus mixed each acid addition with laboratory air, which led to deviations in pH below the set point for both low pCO_2_ treatments. This variability likely resulted from greater headspace pCO_2_ compared to the target level. LCHpH test cultures experienced average decreases of 0.08 ± 0.02 pH units (20.2 ± 4.7% increase in acidity) while LCLpH test cultures saw greater deviations, with an average decrease of 0.21 ± 0.13 pH units (62.2 ± 34.9% increase in acidity). Comparatively, pH in the HCLpH test cultures remained more stable (7.59 ± 0.05 pH units). Revising the apparatus so that cultures were mixed via pCO_2_-matched gas mixtures improved the pH control of each treatment. This refined pH control system consistently maintained pH across all replicate cultures within each treatment group (Tables 1 and S1), where the average pH improved to 8.45 ± 0.02 for the LCHpH treatment, 7.60 ± 0.01 for the HCLpH treatment, and 7.58 ± 0.03 for LCLpH treatment. This comparison to preliminary data is reported to illustrate the difficulty of controlling pH, which will be revisited in the discussion.

Using the revised apparatus, growth rates were similar to maxima for *T. pseudonana* grown at 18°C under 24 h high-intensity light (Guillard and Ryther, 1962; Guillard 1975; Thompson *et al*., 1990; Brown *et al*., 1996; Thompson 1999; Parker and Armbrust, 2005; Laws *et al*., 2020), with observed average growth rates of 1.98 ± 0.07 d^-1^, 2.00 ± 0.08 d^-1^, and 2.03 ± 0.08 d^-1^ for the LCHpH, HCLpH, and LCLpH treatments, respectively (Table 1). Variability in growth rates were similar across experimental conditions (Tables 1 and S2), indicating that the experimental apparatus did not disproportionately affect any one treatment.

**Table 1.** Experimental culture conditions.

| Treatment | Temperature set point (°C) | Target pCO<sub>2</sub> (µatm) | pH<sub>NBS</sub> | Growth rate (d<sup>-1</sup>) |
| --- | --- | --- | --- | --- |
| LCHpH | 18 | 202 | 8.45 ± 0.02 | 1.98 ± 0.07 |
| HCLpH | 18 | 1702 | 7.60 ± 0.01 | 2.00 ± 0.08 |
| LCLpH | 18 | 202 | 7.58 ± 0.03 | 2.03 ± 0.08 |

pH<sub>NBS</sub> and growth rates reported as mean ± SD. For all treatment levels n = 8.

### Proteomic analysis overview

In total, 8,571 proteins were identified with a global false discovery rate of 1%. After quality control filtering, 7,803 accurately quantified proteins remained for further analysis. The comparison of the LCHpH and HCLpH conditions revealed 118 differentially abundant proteins (> 1.30-fold change, adjusted P < 0.05), of which 72 were more abundant while 46 were less abundant in the LCHpH condition compared to the HCLpH condition (Fig. 1 and 2a, Table S3). Twelve differentially abundant proteins were found when comparing the LCHpH and LCLpH conditions, with 6 each more abundant in the LCHpH and LCLpH conditions, respectively (Fig. 2b, Table S4). When identifying differentially abundant proteins common to both low pH treatments, two uncharacterized proteins were found that were more abundant at low pH compared to high pH. Proteins will be referred to by their name and/or Uniprot ID throughout the remainder of this results section.

**Fig. 1.**
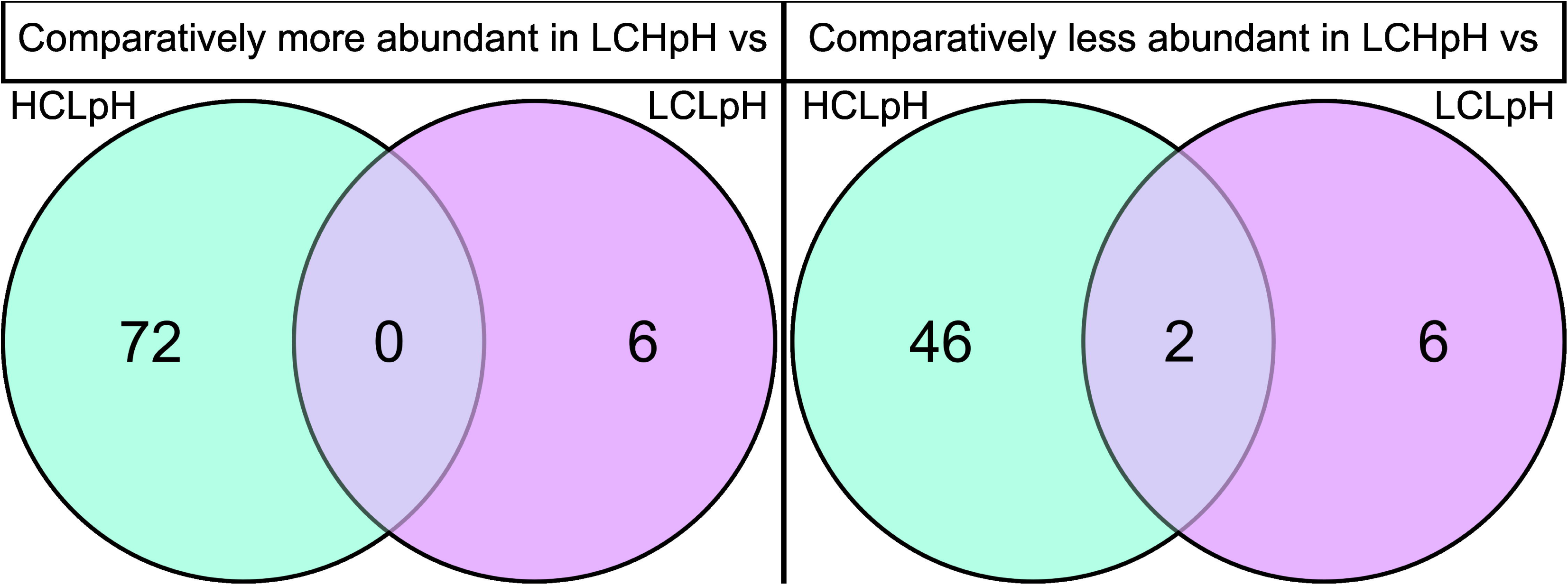
Venn diagrams depicting the number of proteins that were more abundant (left) or less abundant (right) in the LCHpH treatment compared to either the HCLpH (green) or LCLpH (purple) treatments. Overlap indicates differentially expressed proteins shared between both low pH conditions. Graphic generated in Inkscape (Inkscape Project, 2024).

**Fig. 2.**
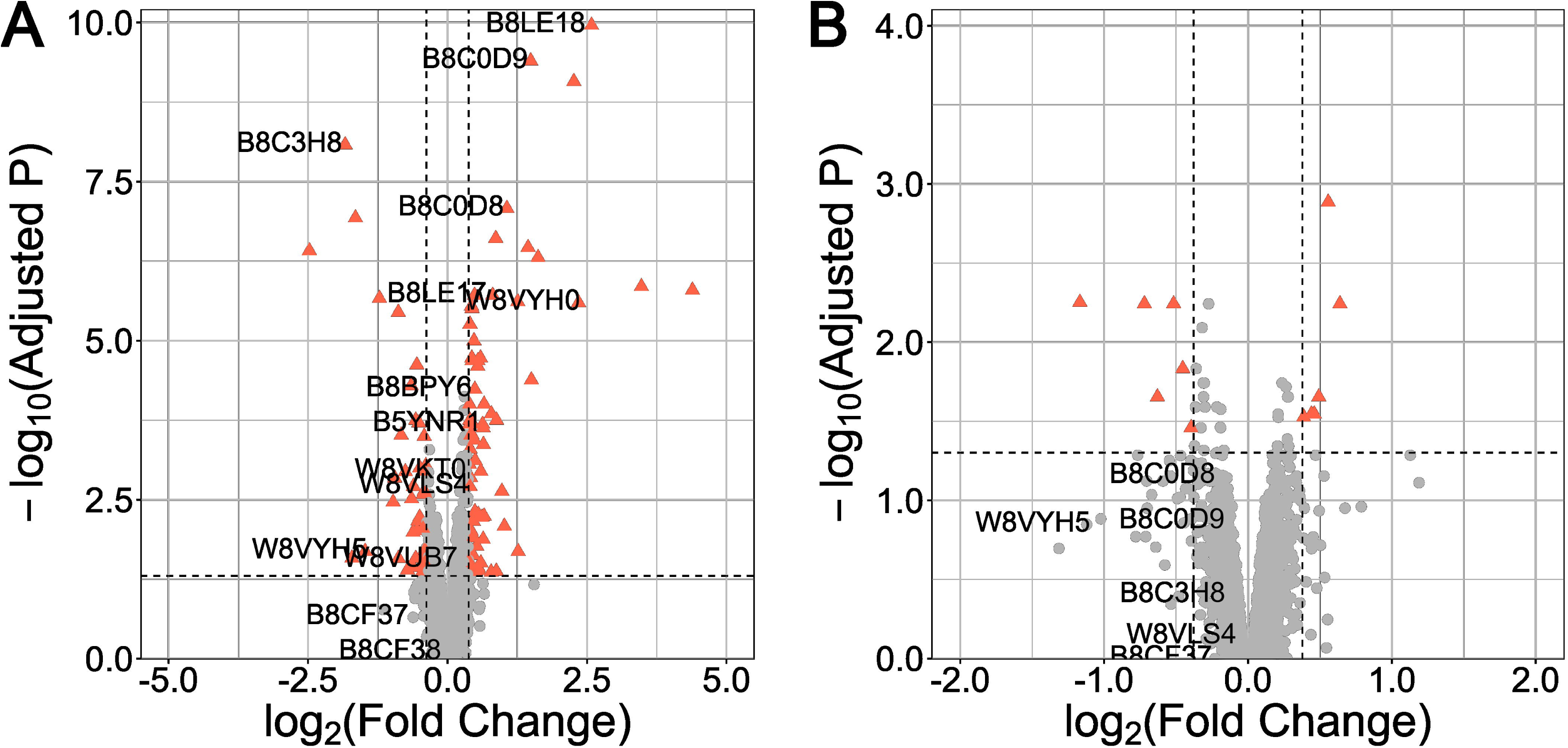
Volcano plots showing changes in protein expression for A) LCHpH compared to HCLpH and B) LCHpH compared to LCLpH. Data shown as -log_10_ of the adjusted P value plotted against log_2_ of the protein abundance fold change. The dashed horizontal line indicates α = 0.05. The dashed vertical lines indicate the *a priori* 1.30 protein fold-change significance threshold. Red triangles represent proteins with significantly different abundances between treatment groups while gray circles represent proteins that did not cross both significance thresholds. The data point markers are shown to the right of the labels, which correspond to Uniprot IDs listed in Table 2. Overlapping labels were removed for readability. Plots produced with ggplot2 R package ver. 3.5.1 (Wickham, 2016).

### Differential abundance of proteins involved in carbon concentrating mechanisms

Seven annotated CAs were differentially abundant between the LCHpH and HCLpH conditions alongside an eighth possible CA (B8LE18; Table 2). δCA1, θCA1, θCA2, θCA3, and ζCA1 were more abundant in cells grown in the LCHpH condition, with respective fold increases of 5.10, 2.39, 1.40, 1.75, and 9.69. αCA3 and θCA4 were less abundant in cells from the LCHpH condition by 1.35- and 3.57-fold, respectively. γCA1, γCA4, and γCAL were also quantified in the LCHpH and HCLpH conditions, and no differences were detected. No differences in the abundance of any CA were found when comparing the low pCO_2_ treatments.

**Table 2.** Differential abundance of key proteins with hypothesized roles in a CCM, pH homeostasis mechanism, cyclic electron transport pathway, and/or excess energy dissipation pathway.

| Uniprot ID | Description | Name | Average fold change<br>LCHpH v HCLpH | Average fold change<br>LCHpH v LCLpH | Source |
| --- | --- | --- | --- | --- | --- |
| B8C0D8 | Bestrophin 1 | TpBST1 | 2.09 | - | Nigishi et al., 2024 |
| B8C0D9 | Bestrophin 2 | TpBST2 | 2.81 | - | Nigishi et al., 2024 |
| B8C0V5 | $\alpha$ carbonic anhydrase 3 | TpαCA3 | -1.35 | - | Samukawa et al., 2014 |
| W8VYH0 | $\delta$ carbonic anhydrase 1 | TpδCA1 | 5.10 | - | Samukawa et al., 2014 |
| W8VSK0 | $\gamma$ carbonic anhydrase 1 | TpγCA1 | - | - | Samukawa et al., 2014 |
| B8C215 | $\gamma$ carbonic anhydrase 4 | TpγCA4 | - | - | Samukawa et al., 2014 |
| W8VLR1 | $\gamma$ carbonic anhydrase like protein | TpγCAL | - | - | Samukawa et al., 2014;<br>Cainzos et al., 2021 |
| B8LE19 | $\theta$ carbonic anhydrase 1 | TpθCA1 | 2.39 | - | Nawaly et al., 2023 |
| B8BPY6 | $\theta$ carbonic anhydrase 2 | TpθCA2 | 1.40 | - | Nawaly et al., 2023 |
| B8LE17 | $\theta$ carbonic anhydrase 3 | TpθCA3 | 1.75 | - | Nawaly et al., 2023 |
| B8C3H8 | $\theta$ carbonic anhydrase 4 | TpθCA4 | -3.57 | - | Nawaly et al., 2023 |
| B8CG97 | $\zeta$ carbonic anhydrase 1 | TpζCA1 | 9.69 | - | Tachibana et al., 2011 |
| B8LE18 | LCIP carbonic anhydrase domain-containing protein |  | 5.98 | - | Armbrust et al., 2004;<br>Bowler et al., 2008 |
| B5YNR1 | Malate dehydrogenase | MDH<sub>c</sub> | 1.55 | - | Armbrust et al., 2004;<br>Bowler et al., 2008 |
| B8CF37 | Mitochondrial alternative oxidase 1 | AOX1 | - | - | Armbrust et al., 2004;<br>Bowler et al., 2008 |
| B8CF38 | Mitochondrial alternative oxidase 2 | AOX2 | - | - | Armbrust et al., 2004;<br>Bowler et al., 2008 |
| B8LBH6 | NADH:ubiquinone reductase (non-electrogenic) |  | - | - | Armbrust et al., 2004;<br>Bowler et al., 2008 |
| W8VUB7 | NAD-dependent malic enzyme | NAD-ME | - | - | Tanaka et al., 2014 |
| B8LBP8 | NAD(P)-binding domain-containing protein |  | 1.33 | - | Armbrust et al., 2004;<br>Bowler et al., 2008 |
| B8C9Q9 | Oxoglutarate/malate translocator | OMT | - | - | Armbrust et al., 2004;<br>Bowler et al., 2008 |
| W8VKT0 | Phosphoenolpyruvate carboxylase 1 | PEPC1 | - | - | Tanaka et al., 2014 |
| W8VLS4 | Phosphoenolpyruvate carboxylase 2 | PEPC2 | 1.32 | - | Tanaka et al., 2014 |
| W8VYH5 | Phosphoenolpyruvate carboxykinase | PEPCK | -2.78 | - | Tanaka et al., 2014 |
| B8CC74 | Plastidial terminal oxidase 1 | PTOX1 | ND | ND | Nawrocki et al., 2015 |
| B8LD44 | Plastidial terminal oxidase 2 | PTOX2 | - | - | Nawrocki et al., 2015 |
| B8C035 | Proton gradient regulation 5 protein | PGR5 | - | - | Zhou et al., 2021 |
| B8BZQ3 | PGR5-like photosynthetic phenotype 1 protein | PGRL1 | - | - | Hertle et al., 2013 |

Uniprot ID numbers provided, but descriptions have been updated with more recent information as cited. Differential abundance (> 1.3 fold-change and adjusted $P < 0.05$ ; Benjamini-Hochberg method) is shown as the average fold change in the LCHpH treatment compared to either the HCLpH or LCLpH treatments ( $n = 4$ for all treatments). Positive fold changes indicate greater protein abundance in LCHpH grown cells while negative values denote lower abundance in LCHpH grown cells. Dashes indicate that no significant difference in abundance was detected. ND indicates the protein was not sufficiently detected for accurate quantification.

In conjunction with CAs, HCO_3_^-^ transport proteins can be important to CCMs as they facilitate the movement of HCO_3_^-^ between subcellular compartments. Two known solute carrier family 4 (SLC4) HCO_3_^-^ transporters, SLC4-1 (B8C6G1) and SLC4-2 (B8BQU6), have been identified in *T. pseudonana* (Nakajima *et al*., 2013), but no differences in their abundances were detected (Tables S5 and S6). Additionally, two anion transporters that may be capable of moving HCO_3_^-^, bestrophin 1 and 2 (Nigishi *et al*., 2024), were more abundant, by 2.09- and 2.81-fold, respectively, in the LCHpH condition compared to the HCLpH condition (Table 2). The abundance of these bestrophins did not differ significantly between the low pCO_2_ treatments.

Regardless of its source, HCO_3_^-^ is required by PEPC to produce OAA. The abundance of PEPC1 did not vary significantly between treatments, although it did show an upward trend in LCHpH cells compared to HCLpH cells (1.26-fold increase, adjusted P < 0.05). PEPC2 was significantly more abundant (1.32-fold) under LCHpH compared to HCLpH conditions, and occurred alongside a 2.78-fold lower abundance of phosphoenolpyruvate carboxykinase (PEPCK), which catalyzes the inverse reaction of PEPC (Table2). While PEPC2 is referred to as the mitochondrial form of PEPC, current predictive localization tools can now provide deeper insight. DeepLoc 2.1 (Ødum *et al*., 2024) predicts a general mitochondrial localization probability of 0.80 with a peripheral mitochondrial membrane association probability of 0.73, suggesting that PEPC2 is associated with the outer surface of mitochondria.

In the model C_4_ plant CCM described above, OAA produced by PEPC is converted to malate by MDH. Three forms of MDH were quantified in this study, but only the abundance of the cytoplasmic form (MDH_c_) varied significantly. The abundance of MDH_c_ (B5YNR1) was 1.55-fold higher in LCHpH cells compared to HCLpH cells (Table 2), with no difference detected between the LCHpH and LCLpH treatments.

The next step in a biochemical CCM would be for the newly formed malate to enter the chloroplast via carboxylate transporters. One such transporter is the 2-oxoglutarate/malate translocator (OMT), the abundance of which was similar between treatment groups. Characterizations of other carboxylate transporters are lacking in *T. pseudonana*, preventing any further evaluation of this aspect of its CCM.

Malate, once inside the chloroplast, can be decarboxylated by ME to release CO_2_ near RuBisCO. The abundance of NAD^+^-dependent ME (NAD-ME, W8VUB7) did not differ significantly between treatments and no other forms of ME have yet been identified in *T. pseudonana*. That said, a NAD(P)-binding domain-containing protein (B8LBP8) that shares its closest identity match (66.3%, Uniprot blast tool) with a short-chain dehydrogenase/reductase (SDR) family oxidoreductase in *Skeletonema marinoi* was 1.33-fold more abundant in cells grown in the LCHpH condition compared to the HCLpH condition. Alternatively, it has been hypothesized that pyruvate carboxylase (PYC) may serve as the decarboxylase in the stroma of *T. pseudonana* (Kustka *et al*., 2014). Of the three known forms of PYC within *T. pseudonana*, only the form with a predicted plastidial localization (Kustka *et al*., 2014; ASAFind 2.0 predicts a chloroplast signal peptide with high confidence) was accurately quantified here. Changes in the abundance of this PYC (B8CE42) did not cross the *a priori* protein fold-change significance threshold for either treatment comparison (Table S5 and S6).

After CO_2_ has been released near RuBisCO, the regeneration of PEP from pyruvate produced during the decarboxylation of malate by ME, or OAA by PYC, would serve to resupply PEPC in the cytoplasm. This conversion can be catalyzed by pyruvate, phosphate dikinase (B8C332) and pyruvate, water dikinase (B8C779); however, neither protein was accurately quantified in this study.

### Differential abundance of proteins in alternative pathways employing the malate/OAA shuttle

The pH homeostasis pathways described by Sakano (1998) rely on several enzymes that may also be involved in the CCM of *T. pseudonana*. The first step described in response to elevated intracellular pH is to increase PEPC activity, which aligns with the observed 1.32-fold increase in PEPC2 abundance in LCHpH cells compared to HCLpH cells. When testing the effect of pH alone, no difference in PEPC2 abundance was detected between the LCHpH and LCLpH conditions. Looking further downstream in this pathway, increased PEPC activity consumes PEP, which removes inhibition on phosphofructokinase allowing glycolytic reactions to proceed that yield net H^+^ ions. No significant differences in abundance were detected in any form of phosphofructokinase and furthermore, no differences were observed in the five detected forms of pyruvate kinase, an enzyme also capable of consuming PEP (Table S5 and S6).

The complementary side of this pH homeostasis mechanism that functions as a proton sink relies on NAD-ME to supply NADH + H^+^ into the alternative respiratory pathway. No differences were detected among the three treatments for the only known form of NAD-ME (W8VUB7) in *T. pseudonana*. Downstream of ME, this pH regulatory pathway is dependent upon mitochondrial alternative oxidases (AOX) to transfer protons from NADH + H^+^ to molecular oxygen, reducing it to water (see reviews by Selinski and Scheibe, 2018; Chadee *et al*., 2021). No significant differences in the abundance of either AOX1 (B8CF37) or AOX2 (B8CF38) were found (Table 2). A non-electrogenic form of mitochondrial electron transport chain complex I (B8LBH6), which when paired with AOX can facilitate the oxidation of NADH + H^+^ without altering cellular energy balance, was also detected and no significant differences in its abundance were observed. Although not described in Sakano (1998), the alternative oxidases within the chloroplast, which function similarly to AOX while utilizing NADPH + H^+^, could also act as a proton sink when working in conjunction with the malate/OAA shuttle. While plastidial terminal oxidase 1 (PTOX1, B8CC74) could not be quantified, no variations in the abundance of PTOX2 (B8LD44) were observed (Table 2).

Another cellular mechanism that can involve the malate/OAA shuttle is cyclic electron transport around photosystem I (CET; reviewed in Chadee *et al*., 2021). The two known pathways of CET are dependent on 1) the proton gradient regulation 5 protein (PGR5, B8C035) in conjunction with the PGR5-like photosynthetic phenotype 1 protein (PGRL1, B8BZQ3) and 2) the NADH-like (NDH) complex. No significant differences in the abundances of PGR5 and PGRL1 were detected (Table 2), and while numerous subunits are known to comprise the NDH complex in *T. pseudonana*, many remain poorly characterized making its evaluation difficult.

## Discussion

The proteomic analysis resulted in 7,803 quantified proteins following rigorous filtering. This represents one of the most complete *T. pseudonana* proteomes to date, and therefore, allowed for an in-depth exploration of both the CCM and other metabolic pathways that could produce similar protein abundance profiles. Differential protein abundances are discussed in a stepwise fashion proceeding through the proposed CCM of *T. pseudonana* (Fig. 3). This is followed by a discussion of intracellular CO_2_ recapture (Fig. 4) before alternative explanations for increased abundances of C_4_ CCM enzymes are evaluated. This section concludes with an examination of challenges inherent to exploring the CCM of *T. pseudonana* and strategies to avoid them moving forward.

**Fig. 3.**
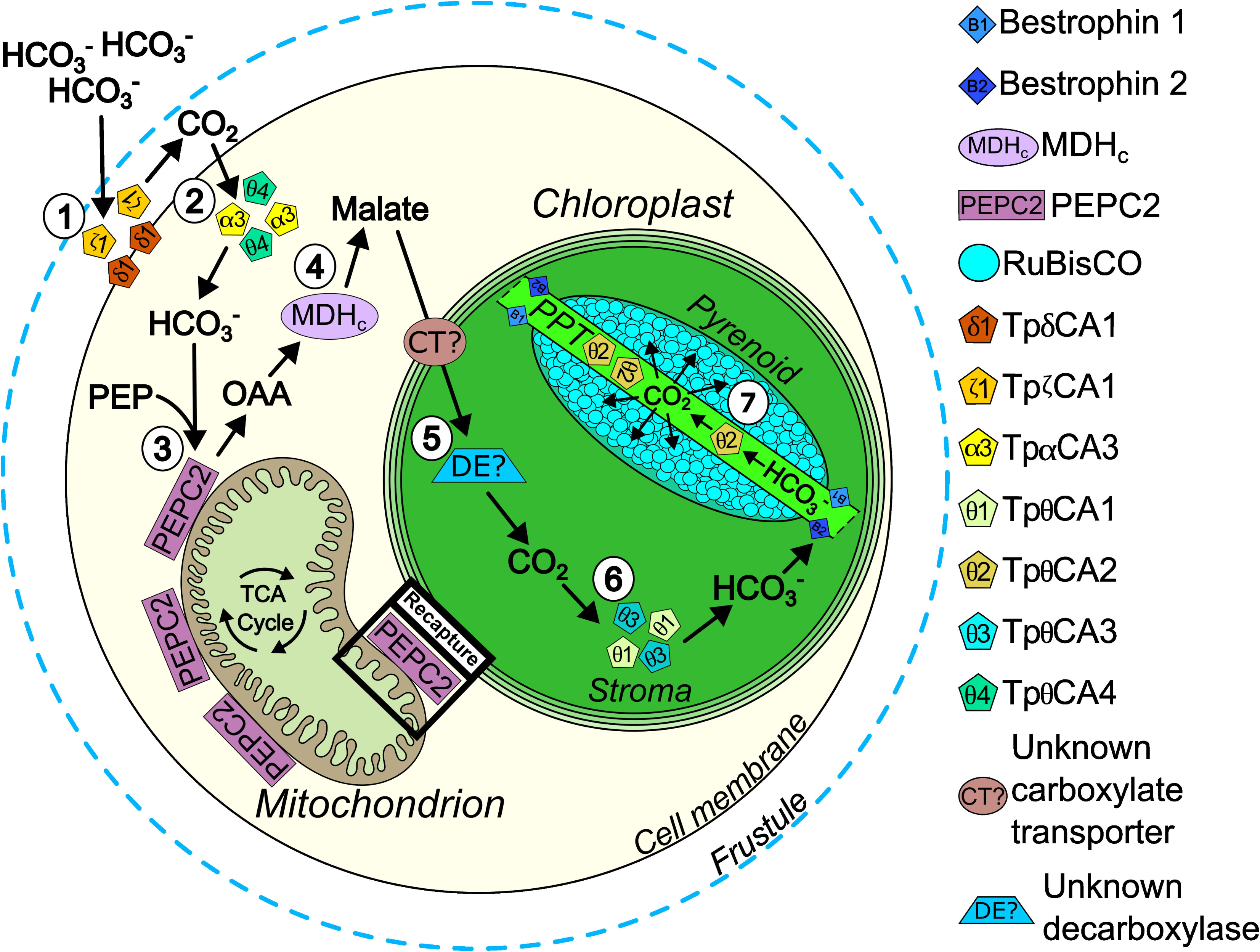
Cross section of a *T. pseudonana* cell showing the schematic of the proposed CCM. Concentric circles starting from the outside moving in represent the frustule, plasma membrane, and the four membranes of the complex plastid. Each step of the CCM is numbered following the subheadings within the discussion. **Step 1:** the conversion of extracellular HCO_3_^-^ to CO_2_ by TpδCA1 and TpζCA1 which then diffuses across the plasma membrane. **Step 2:** the conversion of this influxing CO_2_ back to HCO_3_^-^ potentially by TpαCA3 and TpθCA4. **Step 3:** the carboxylation of PEP and HCO_3_^-^ to OAA by PEPC2 (or the class 2 PEPC complex). **Step 4:** the conversion of this OAA to malate by MDH_c_. **Step 5:** the translocation of malate into the chloroplast by an as yet unknown carboxylate transporter before the unknown stromal decarboxylase releases CO_2_ from malate. **Step 6:** the conversion of CO_2_ produced by the decarboxylase into HCO_3_^-^ by TpθCA1 and TpθCA3 before HCO_3_^-^ then diffuses down its concentration gradient through bestrophin 1 and 2 into the PPT. **Step 7:** TpθCA2 within the PPT lumen converts influxing HCO_3_^-^ back to CO_2_, which then passively diffuses out to RuBisCO in the pyrenoid. The black box titled recapture across the mitochondrial membranes corresponds to the zoomed in view of the proposed CO_2_ recapture mechanism shown in Figure 4. Cellular components are not shown to scale. Abbreviations ordered stepwise: δ1, delta-type carbonic anhydrase 1; ζ1, zeta-type carbonic anhydrase 1; α3, alpha-type carbonic anhydrase 3; θ4, theta- type carbonic anhydrase 4; PEP, phosphoenolpyruvate; PEPC2, phosphoenolpyruvate carboxylase 2; OAA, oxaloacetate; MDH_c_, cytoplasmic form of malate dehydrogenase; CT?, unknown carboxylate transporter; DE?, unknown decarboxylase; θ1, theta-type carbonic anhydrase 1; θ3, theta-type carbonic anhydrase 3; B1, bestrophin 1; B2, bestrophin 2; PPT, pyrenoid-penetrating thylakoid; θ2, theta-type carbonic anhydrase 2. Graphic generated in Inkscape (Inkscape Project, 2024).

**Fig. 4.**
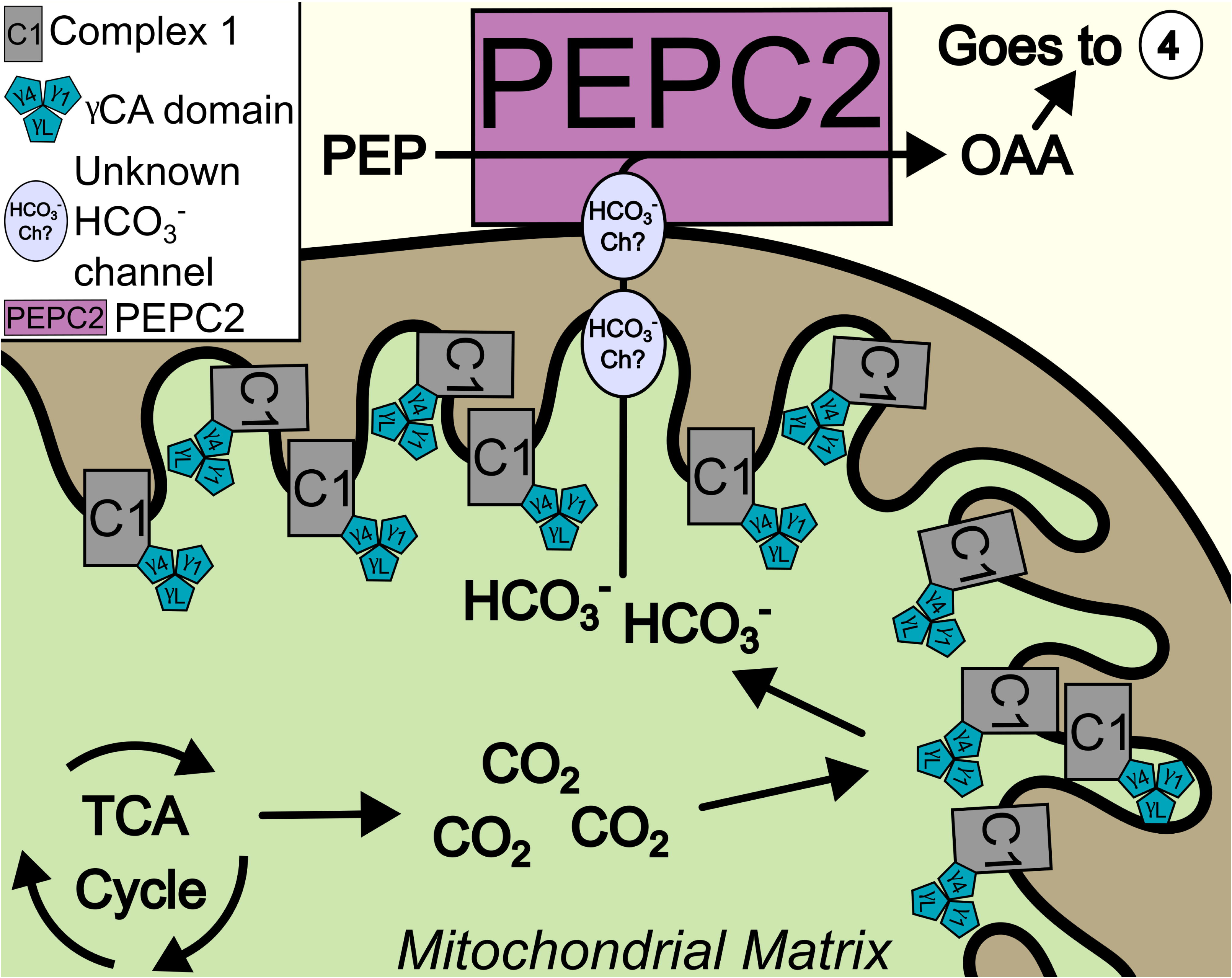
Magnified view of a portion of a mitochondrion within *T. pseudonana* showing the proposed CO_2_ recapture mechanism. CO_2_ is produced as a byproduct of the tricarboxylic acid (TCA) cycle in the mitochondrial matrix and begins to diffuse out of the mitochondrion. Once CO_2_ reaches the inner mitochondrial membrane it encounters the γCA domain, which is composed of three heterogeneous subunits, within a portion of complex I that extends into the matrix. The γCA domain converts CO_2_ to HCO_3_^-^ which leaves the mitochondrion through unknown HCO_3_^-^ channels out to PEPC2 (or the class 2 PEPC complex). PEPC2 then converts this HCO_3_^-^ and PEP into OAA, which continues on into step 4 of the CCM shown in Figure 3. Cellular components are not shown to scale. Abbreviations: C1, mitochondrial electron transport chain complex 1; γ1, gamma-type carbonic anhydrase 1; γ4, gamma-type carbonic anhydrase 4; γL, gamma-type carbonic anhydrase like protein; HCO_3_^-^ Ch?, unknown bicarbonate channel; PEP, phosphoenolpyruvate; PEPC2, phosphoenolpyruvate carboxylase 2; OAA, oxaloacetate. Graphic generated in Inkscape (Inkscape Project, 2024).

### Carbon concentrating mechanism

#### Step 1: Extracellular DIC uptake

CCMs must begin with the uptake of extracellular DIC in the form of CO_2_ or HCO_3_^-^, which can be facilitated by externally acting CAs and HCO_3_^-^ transporters (reviewed by Reinfelder *et al*., 2011). TpδCA1 and TpζCA1 are both believed to operate in the periplasmic space (Tachibana *et al*., 2011; Samukawa *et al*., 2014; Hopkinson *et al*., 2016) and showed two of the largest increases in protein abundance in LCHpH compared to HCLpH conditions (Table S3). The upregulation of external CAs to produce HCO_3_^-^ would seemingly be of little benefit if HCO_3_^-^ uptake is the initial step of the CCM, as the total DIC pool at both pH 7.61 and 8.48 is predominantly HCO_3_^-^ (Mehrbach *et al*., 1973; Dickson and Millero, 1987; Lueker *et al*., 2000). Furthermore, recent work has shown that *T. pseudonana* is insensitive to a known inhibitor of extracellular SLC4 family HCO_3_^-^ transporters while exposure to an extracellular CA inhibitor produced a marked decrease in its affinity for DIC (Tsuji *et al*., 2017). The current study found no significant changes in the abundance of any potential extracellular HCO_3_^-^ transporter or ATPase, at least a pair of which would be needed to facilitate increased HCO_3_^-^ uptake (Thielmann *et al*., 1990; Karlsson *et al*., 1994; Nakajima *et al*., 2013; Wang *et al*., 2015; Poschenrieder *et al*., 2018; Nawaly *et al*., 2023a). Taken together, these results further solidify the first step of the CCM in *T. pseudonana* as the passive influx of CO_2_ generated by external CA activity (Moroney *et al*., 2011; Hopkinson *et al*., 2013; Kustka *et al*., 2014; Tsuji *et al*., 2017; Tsuji *et al*., 2021).

### Step 2: The role of cytoplasmic carbonic anhydrases

The next logical step would be the “capture” of CO_2_ entering the cell through its CA-mediated conversion back to HCO_3_^-^ to prevent diffusive losses. Two cytosolic forms of CA, TpαCA3 and TpθCA4 (Tachibana *et al*., 2011; Nawaly *et al*., 2023b), were differentially abundant in this study; however, they were more abundant in the HCLpH compared to the LCHpH condition. As CAs can serve dual roles to both concentrate DIC and regulate cellular acid-base status, contrasting interpretations must be considered (Reinfelder *et al*., 2011; Supuran, 2016; Occhipinti and Boron, 2019). Increased abundances of TpαCA3 and TpθCA4 at high pCO_2_ could indicate that these CAs act to buffer cytoplasmic acidification driven by the greater influx of CO_2_ (Badger and Price, 1994; Occhipinti and Boron, 2019). This would not preclude them from converting influxing CO_2_ to HCO_3_^-^ under low pCO_2_ conditions, but instead could simply indicate that they are needed in greater abundance for acid-base regulation at a pCO_2_ of 1702 μatm compared to that needed for CO_2_ capture at a pCO_2_ of 202 μatm. Conversely, decreased abundances of TpαCA3 and TpθCA4 in response to low pCO_2_ would negate their involvement in the CCM. The totality of low pCO_2_-induced changes in protein abundances when considered with the lack of corresponding changes in HCO_3_^-^ uptake mechanisms; however, suggest that *T. pseudonana* relies on cytoplasmic CAs to convert influxing CO_2_ to HCO_3_^-^ at both tested pCO_2_ levels.

### Step 3: OAA formation

PEPC2 abundance in *T. pseudonana* increased in response to low pCO_2_ but not extracellular pH, indicating that it could serves as a C_4_ carboxylase in the CCM. As current predictions for *T. pseudonana*, which concur with observations in other phototrophs (O’Leary *et al*., 2011; Park *et al*., 2012; O’Leary and Plaxton, 2020), place PEPC2 on the peripheral surface of the mitochondria, it likely utilizes cytoplasmic pools of PEP and HCO_3_^-^ to produce OAA. This localization still allows PEPC2 to serve in its anaplerotic role for the TCA cycle through coupling with known mitochondrial dicarboxylate transporters (O’Leary *et al*., 2011; Park *et al*., 2012; Schober *et al*., 2019; Fernie *et al*., 2020; Liu *et al*., 2022; Zara *et al*., 2022) while also preventing the futile cycling of OAA through the spatial isolation of PEPC2 and PEPCK (Kustka *et al*., 2014). The observed decrease in PEPCK abundance under low pCO_2_ conditions would further support maximal CCM functioning by reducing OAA demand within the mitochondria. While more work is needed to confirm the localization of PEPC2, there is another discovery pertaining to both PEPC1 and PEPC2 that could have major implications for past findings on the role of PEPC in the CCM of *T. pseudonana*.

Work in green algae and land plants has revealed the existence of a heteromeric PEPC complex (termed the class 2 PEPC complex), composed of four subunits each of PEPC1 and PEPC2, that may be the only form in which PEPC2 subunits function within these taxa (Rivoal *et al*., 1996; Rivoal *et al*., 1998; Rivoal *et al*., 2002; Mamedov *et al*., 2005; Moellering *et al*., 2007; O’Leary *et al*., 2011; Park *et al*., 2012; Ting *et al*., 2017; Koendjbiharie *et al*., 2021; Hu *et al*., 2025). This complex is localized to the surface of the mitochondria and is less sensitive, relative to homotetrameric PEPC1 complexes, to allosteric inhibition by metabolic intermediates, such as malate and aspartate (O’Leary *et al*., 2009; O’Leary *et al*., 2011; Park *et al*., 2012; Ting *et al*., 2017). As will be discussed more fully below, malate likely plays a central role in the biochemical aspects of the CCM in *T. pseudonana*, and therefore, the presence of a malate- resistant PEPC complex would help to facilitate optimal CCM functioning. If the presence of a class 2 PEPC complex in *T. pseudonana* is verified, it would call for a reexamination of previously reported PEPC data. Parallel increases in both PEPC1 and PEPC2 (Granum *et al*., 2009; Kustka et al 2014) could be indicative of the upregulation of the hybrid class 2 complex. While the change observed here in PEPC1 abundance did not cross the *a priori* protein fold change significance threshold, the 1.26-fold (adjusted P = 0.001) increase under LCHpH compared to HCLpH conditions is similar to the 1.32-fold increase in PEPC2 abundance, which would align with increased demand for the class 2 complex. Additionally, careful consideration of PEPC activity measurements would be warranted, as active class 2 complexes have proven difficult to extract given their high susceptibility to cleavage by thiol endopeptidases (Blonde and Plaxton, 2003; Gennidakis *et al*., 2007; O’Leary *et al*., 2009). Although this remains speculative, it highlights the need for future work exploring the molecular structure of diatom PEPC complexes in order to fully understand the operation and regulation of their CCMs.

### Step 4: Malate formation and transport

The abundance of MDH_c_ increased in response to low pCO_2_ conditions, but was unaffected by extracellular pH, suggesting that OAA is reduced to malate in the cytosol before it is translocated into the chloroplast stroma. Malate exchange across the chloroplast envelope can readily occur as several plastidial di- and tri- carboxylate transporters (e.g. OMT, DiT2) are less substrate specific than initially believed (Taniguchi *et al*., 2002; Kinoshita *et al*., 2011; Selinski and Scheibe, 2018). While no change in the abundance of OMT was detected, and no increase in any transporter may be necessary if malate translocation occurs rapidly, more work is needed to characterize other plastidial carboxylate transporters in *T. pseudonana* as few are well described.

### Step 5: Decarboxylation near RuBisCO

In the simplest C_4_ plant CCM, NADP^+^-dependent malic enzyme (NADP-ME) decarboxylates malate back to pyruvate while releasing CO_2_ (reviewed by Leegood, 2002). The only annotated ME in *T. pseudonana* showed similar abundances across all treatment groups in this study and to date, no forms of NADP-ME or additional forms of NAD-ME have been observed (Tanaka *et al*., 2014; Davis *et al*., 2017). However, an NAD(P)-binding domain-containing protein (B8LBP8) was significantly more abundant in cells grown under low pCO_2_ (Table 2). Uniprot blast tool results showed that its closest identity match is to an SDR family NAD(P)-dependent oxidoreductase (A0AAD8Y4P9) found in another centric diatom, *S. marinoi*. While the SDR family of oxidoreductases contains thousands of proteins, it is the family within which ME resides (Kavanagh *et al*., 2008; Moon *et al*., 2012; Tronconi *et al*., 2018). The NAD(P)-binding domain of B8LBP8 encapsulates all but 29 of its amino acids with no other known domains detected (NCBI conserved domain search tool, Marchler-Bauer *et al*., 2017; Lu *et al*., 2020; Wang *et al*., 2023), indicating that this protein sequence may be incomplete. While this by no means identifies the decarboxylase in the CCM of *T. pseudonana*, it does provide a direction for future research into uncharacterized oxidoreductases that may not be fully represented in the current reference genome.

Alternatively, it has been proposed that PYC could serve as the final decarboxylase within *T. pseudonana* (Kustka *et al*., 2014). In this scenario, OAA is transported into the stroma instead of malate where its subsequent decarboxylation by PYC releases CO_2_. The only form of PYC detected in this analysis has a predicated localization to the chloroplast, and while there was an increasing trend in its abundance in the LCHpH condition compared to the HCLpH condition, this was not significant. Given the previously reported increase in this PYC under low CO_2_ conditions (Kustka *et al*., 2014) and the two forms of PYC that could not be quantified here, a more targeted assessment of the role of PYC in the CCM of *T. pseudonana* may be warranted. However, if PYC is the stromal decarboxylase, the role of MDH_c_ would need to be reevaluated as it would seemingly limit CCM functioning by reducing the availability of cytosolic OAA (reviewed in Selinski and Scheibe, 2018).

### Step 6: Crossing the pyrenoid protein shell

Recent advancements localizing the θCAs and resolving the pyrenoid ultrastructure in *T. pseudonana* have helped to elucidate the most likely final steps of the CCM within the chloroplast (Nawaly *et al*., 2023b; Nam *et al*., 2024; Shimakawa *et al*., 2024). The abundances of two stromal θCAs, TpθCA1 and TpθCA3, increased in response to low pCO_2_ conditions, but were unaffected by extracellular pH. It has been hypothesized that one or both of these CAs act to recapture CO_2_ that escapes the pyrenoid-penetrating thylakoid (PPT; Nawaly *et al*., 2023b); however, here it is proposed that they primarily serve to convert CO_2_ released from the decarboxylation of malate into HCO_3_^-^. Morphological similarities between prokaryotic carboxysomes and the pyrenoid protein shell of *T. pseudonana* suggest that the pyrenoid could be impermeable to CO_2_ (Nam *et al*., 2024; Shimakawa *et al*., 2024), which would reduce the need for a dedicated CO_2_ recapture mechanism in the stroma. Once TpθCA1 and TpθCA3 capture CO_2_ released from the decarboxylation of malate, the resultant HCO_3_^-^ needs to pass into the PPT lumen where the final step of the CCM resides. Bestrophin 1 and 2, which have recently been localized to the PPT membrane in *T. pseudonana* (Nigishi *et al*., 2024), could facilitate this movement as bestrophins are known to be permeable to HCO_3_^-^ (Qu and Hartzell, 2008). Furthermore, the passage of HCO_3_^-^ through bestrophin channels into the PPT lumen has been documented in several other marine algae (Mukherjee *et al*., 2019; Nigishi *et al*., 2024). This function is similarly supported in *T. pseudonana* as the abundances of bestrophin 1 and 2 increased in response to low pCO_2_ in both this study and others (Kustka *et al*., 2014; Nigishi *et al*., 2024).

### Step 7: Dehydration of HCO_3_^-^

TpθCA2 has been localized within the PPT, where it likely converts stroma-derived HCO_3_^-^ to CO_2_ that then diffuses to RuBisCO in the interior of the pyrenoid (Nawaly *et al*., 2023b; Nam *et al*., 2024; Shimakawa *et al*., 2024). This role is supported here as TpθCA2 abundance increased in response to low pCO_2_, but was insensitive to changes in external pH. While several key areas still require additional characterization, taken as a whole, the results presented here are consistent with a hybrid CCM in *T. pseudonana* that consists of both biophysical and biochemical steps working in unison to supply CO_2_ to RuBisCO.

### Intracellular CO_2_ recapture mechanism

Several CA-mediated recapture mechanisms of respired CO_2_ have been proposed for *T. pseudonana* (Trimborn *et al*., 2009; Tachibana *et al*., 2011; Samukawa *et al*., 2014; Nawaly *et al*., 2023b). However, the strongest case for recycling involves γCA-mediated recapture of CO_2_ from the TCA cycle (Fig. 4; Cardol *et al*., 2004; Parisi *et al*., 2004; Mitra *et al*., 2005; Perales *et al*., 2005; Sunderhaus *et al*., 2006; Braun and Zabaleta, 2007; Zabaleta *et al*., 2012; Soto *et al*., 2015; Fromm *et al*., 2016a; Fromm *et al*., 2016b; Cainzos *et al*., 2021; Klusch *et al*., 2023; Maldonado *et al*., 2023; Braun and Klusch, 2024). Structural analyses in other phototrophs have shown that γCAs and γCA-like proteins (γCAL) form a domain within complex I of the mitochondrial electron transport chain (ETC) that protrudes into the mitochondrial matrix (Sunderhaus *et al*., 2006; Meyer *et al*., 2011; Klusch *et al*., 2023; Maldonado *et al*., 2023) where it converts CO_2_ to HCO_3_^-^. This γCA domain has not been found to function freely of complex I (Martínez-Reyes and Chandel, 2020; Cainzos *et al*., 2021; Klusch *et al*., 2023; Maldonado *et al*., 2023; Braun and Klusch, 2024), indicating that the cellular demand for complex I likely dictates its expression. Recent work in *T. pseudonana* has shown that γCA1, γCA4, and γCAL are capable of forming a heterotrimeric domain with a functional active site (Cainzos *et al*., 2021). The abundance profiles of these γCA subunits and other complex I components were similar among the treatment groups assessed here, which is consistent with this mechanism. A mitochondrial CA-dependent recapture mechanism has been previously considered in algae and was suggested to be energetically favorable. Specifically, Raven (2001) hypothesized that HCO_3_^-^ channels could exist in the mitochondrial membranes of algae to move recaptured DIC without the risk of CO_2_ leakage. It was estimated that these HCO_3_^-^ channels could increase the supply of CO_2_ to RuBisCO by 10% at a cost of <1% of total cellular ATP production. To date, few HCO_3_^-^ channels are known in *T. pseudonana* (Nakajima *et al*., 2013; Tsuji *et al*., 2017) and localization studies are lacking, so more work is needed to further evaluate this recapture mechanism.

### Evidence for alternative cellular pathways employing similar machinery

The quantitative proteomic analysis of *T. pseudonana* allowed for the evaluation of whether the observed upregulation of PEPC and MDH_c_ were driven by demand for the CCM or instead by increased demand for the transport of reducing equivalents via the malate/OAA shuttle. This shuttle in conjunction with the alternative oxidases, which act as shunts that uncouple electron transport to preserve redox poise at the cost of ATP production, can be involved in pH homeostasis, cyclic electron transport, and excess energy dissipation (Shikanai, 2007; Yoshida *et al*., 2007; van Dongen *et al*., 2011; Yu *et al*., 2014; Nawrocki *et al*., 2015; Dietz *et al*., 2016; Murik *et al*., 2018; Chadee *et al*., 2021; Zhou *et al*., 2021; Hippmann *et al*., 2022; Degen *et al*., 2024). Studies have shown that the alternative oxidases are active in cellular functions ranging from respiration to stress tolerance, with correspondingly complex regulatory pathways tightly controlling their expression (Yoshida *et al*., 2007; van Dongen *et al*., 2011; Nawrocki *et al*., 2015; Murik *et al*., 2018; Vanlerberghe *et al*., 2020).

When evaluating the possible involvement of the previously described pH homeostasis pathways (Sakano, 1998), the lack of changes in the abundances of both AOX and PTOX between either low pH treatment and the high pH treatment indicates that the pathway to reduce intracellular H^+^ concentration was not upregulated at an extracellular pH of 7.61. There was also no sign that the H^+^ generating pathway was active in cultures grown at pH 8.48 as no changes in PEPC abundance were detected between the LCHpH and LCLpH groups.

The malate/OAA shuttle is also known to support some forms of cyclic electron transport (CET) around photosystem I (Shikanai, 2007; Nawrocki *et al*., 2015; Chadee *et al*., 2021). CET on its own circumvents NADPH + H^+^ production while still producing ATP to help correct energy imbalances caused by increased ATP consumption. When paired with either PTOX or AOX via the malate/OAA shuttle, CET aids in NADP^+^ regeneration, thereby maintaining both energy balance and redox poise within the chloroplast (Yamori *et al*., 2015; Chadee *et al*., 2021). Of the two known CET pathways, the PGR5 pathway and the NDH pathway (Yamori *et al*., 2015; Chadee *et al*., 2021; Zhou *et al*., 2021; Degen *et al*., 2024), PGR5-dependent CET is more well characterized in *T. pseudonana*. No proteins uniquely involved in this pathway were differentially abundant in this study, and while the NDH pathway is not fully characterized in *T. pseudonana*, there was no indication of differential abundance of any suspected NDH complex subunit (Tables S5 and S6). When considered in conjunction with the lack of changes in AOX and PTOX, there is no evidence that CET is responsible for the results obtained in this study.

Lastly, a mechanism for dissipating excess light energy while combating oxidative stress that relies on the non-electrogenic homolog of mitochondrial complex I, the malate/OAA shuttle, and AOX was examined (Selinski and Scheibe, 2018). In addition to the lack of differences in the AOXs and the non-electrogenic form of complex I, no indications of elevated oxidative stress (e.g. more abundant superoxidase dismutase, catalase, glutathione peroxidase, glutathione reductase; Tables S5 and S6) were observed, demonstrating that this energy dissipation mechanism is not differentially active among treatments. Overall, no evidence of any alternative mechanism was found that could detract from the conclusion that PEPC and MDH_c_ are active in the CCM of *T. pseudonana*.

### Experimental challenges and strategies moving forward

Achieving stringent pH control in all treatments required extensive testing and a refined experimental apparatus. pH control is critical as *T. pseudonana* exhibits rapid changes in carbon metabolism in response to changes in pH and pCO_2_. (Roberts *et al*., 2007; Granum *et al*., 2009; Kustka *et al*., 2014; Clement *et al*., 2016; Clement *et al*., 2017). That said, some *T. pseudonana* CCM studies provide little to no quantification of the carbonate chemistry within their experimental cultures (Table 3), making true comparisons between them challenging. Maintaining consistent carbonate chemistry conditions has proven difficult when diatom cultures are grown to high biomass (Shi *et al*., 2009; Crawfurd *et al*., 2011); however, the specific design of an experimental apparatus can also lead to poorly controlled carbonate chemistry at even low biomass, as was reported here for the preliminary experimental setup. While the lack of carbonate chemistry monitoring does not necessarily indicate that culture conditions were poorly maintained, large deviations from well-established growth rates are more suggestive of the influence of unknown factors.

**Table 3.**
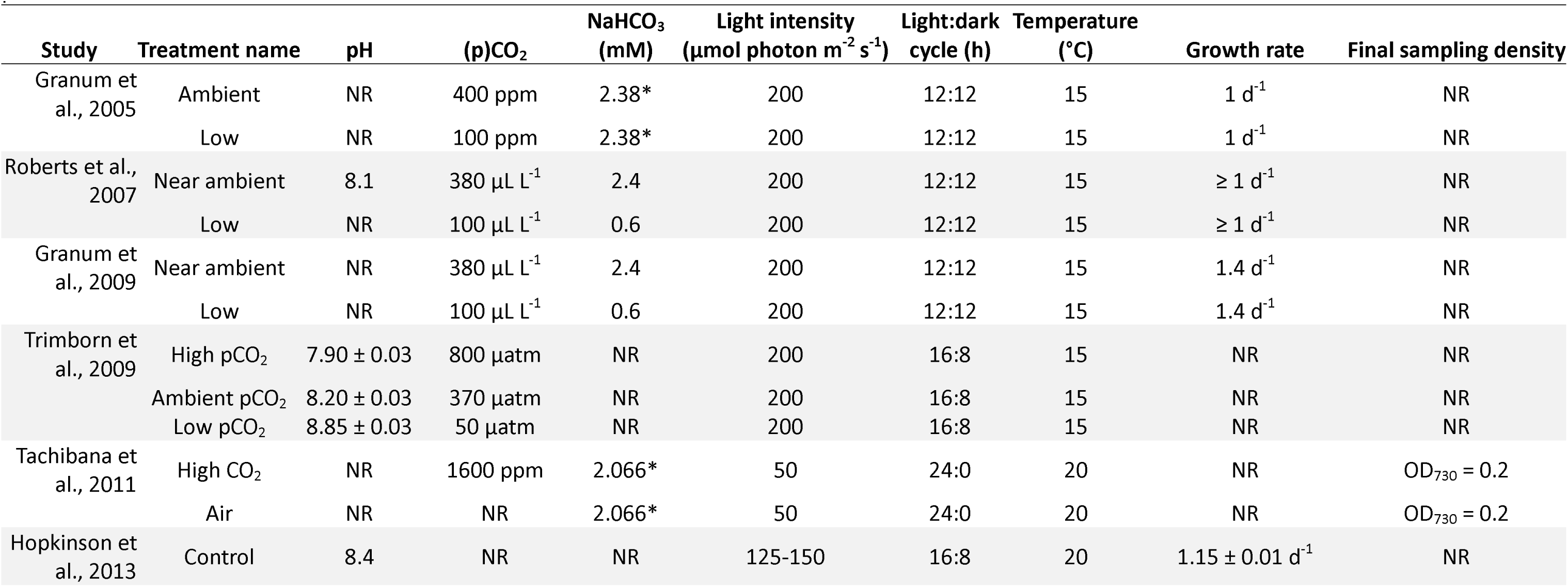
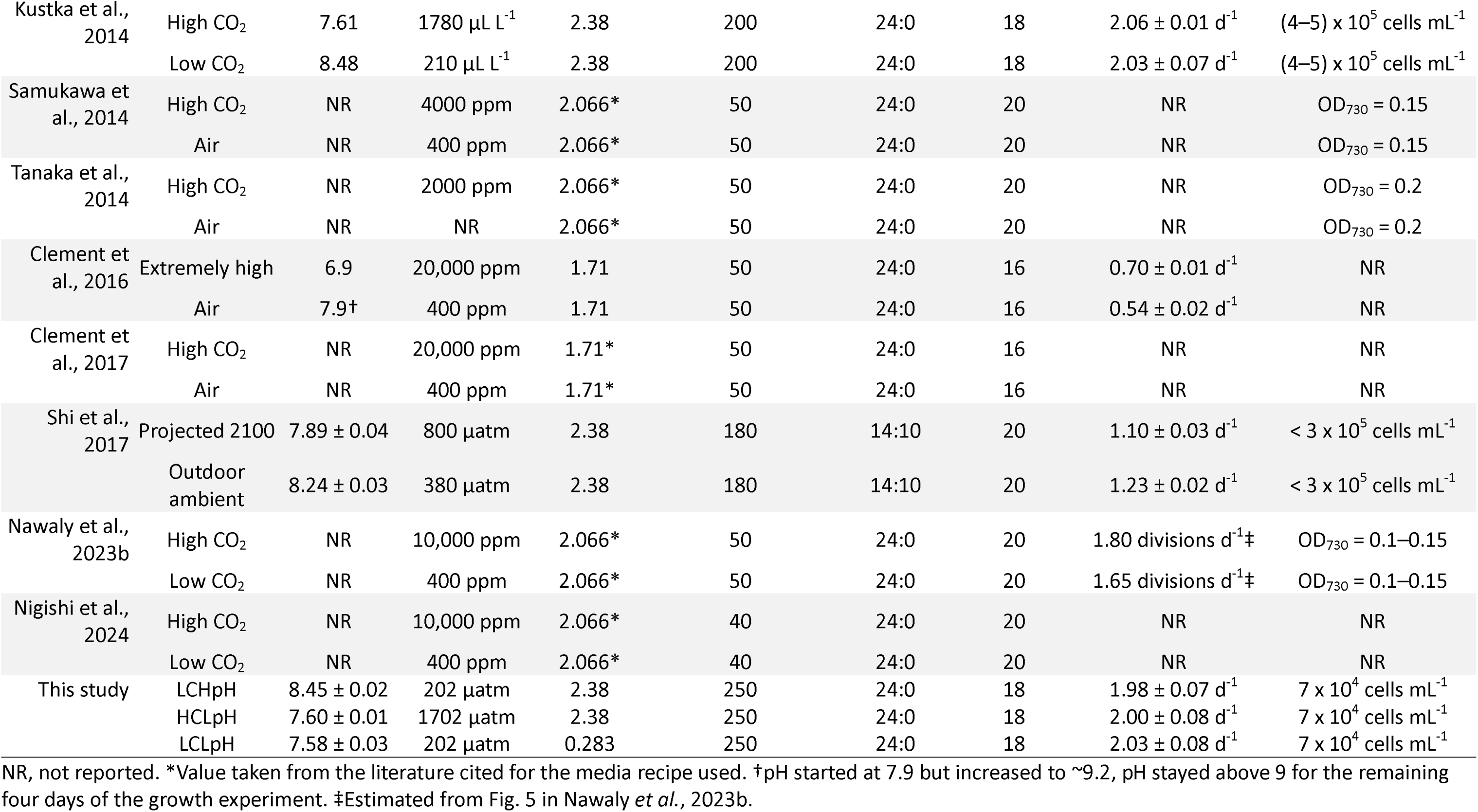
Steady state culture conditions, growth rates, and final culture sampling densities reported from a selection of *T. pseudonana* CCM studies cited within the present work.

As growth is a widely used metric to assess the health of algal cultures, many studies have measured growth rates of *T. pseudonana* (CCMP 1335; previously clone 3H) across a range of temperatures, light conditions, and nutrient concentrations (Guillard and Ryther, 1962; Guillard *et al*., 1973; Guillard, 1975; Thompson *et al*., 1990; Brown *et al*., 1996; Thompson 1999; Parker and Armbrust, 2005; Shi *et al*., 2015; Laws *et al*., 2020). This body of work has shown that nutrient replete *T. pseudonana* cultures grown between 15 and 20°C with continuous low-intensity light (∼50 μmol photon m^-2^ s^-1^) should exhibit a minimum growth rate of approximately 1.5 d^-1^ (2.2 divisions d^-1^). Similar cultures grown with higher intensity light (> 150 μmol photon m^-2^ s^-1^) should achieve growth rates from 1.8 d^-1^ to as high as 2.3 d^-1^ (2.6 and 3.3 divisions d^-1^, respectively). While some variability is expected across studies, large deviations from these values are likely indicative of confounding variables adversely affecting *T. pseudonana*. Studies reporting lower than expected growth rates as well as those where growth is not reported are common in the *T. pseudonana* CCM literature (Table 3). Comparing studies with growth rates that deviate to varying degrees from well characterized expectations can lead to inadvertently erroneous conclusions. To ensure new findings are properly contextualized, all physiological studies should report calculated growth rates moving forward.

In the present study, high and low pCO_2_ *T. pseudonana* cultures were maintained under conditions that yielded maximal growth rates (Guillard and Ryther, 1962; Thompson 1999; Parker and Armbrust, 2005; Laws *et al*., 2020). As cellular carbon demand is predicated on growth, maximally growing low pCO_2_ cultures would have the greatest need for elevated CCM expression. This approach was implemented over the use of extreme CO_2_ levels (from 50 to 20,000 ppm; Clement *et al*., 2016; Clement *et al*., 2017; Jensen *et al*., 2019; Nawaly *et al*., 2023b; Nigishi *et al*., 2024) as using levels that far exceed those experienced in the evolutionary history of diatoms could produce physiological responses unrelated to the CCM (Kuhn and Kasting, 1983; Kasting, 1993; Kah and Riding, 2007; Feulner, 2012; Novák Vanclová and Dorrell, 2024). To accurately interpret results generated using extreme CO_2_ conditions, evidence from additional intermediate CO_2_ levels is needed to ensure observed responses are representative of CCM functioning alone. It is therefore suggested that future studies employ more reasonable pCO_2_ levels to alleviate the need for expansive factorial designs and to provide added insights into how this species will respond to projected future changes in ocean carbonate chemistry.

## Conclusion

The proteomic data presented in this study demonstrate strong evidence for a hybrid CCM within *T. pseudonana*. These results agree with recent work that both the initial and final steps of the CCM are biophysical, while the intermediate steps rely on C_4_ intermediates to translocate CO_2_ near to RuBisCO. Future investigations into the identity and structure of the PEPC complexes present within *T. pseudonana*, the identity of the stromal decarboxylase, and the potential presence of mitochondrial HCO_3_^-^ channels will provide clarity on the as yet unelucidated steps within this CCM. The proteomic approach used here also provided the first in-depth evaluation of alternative pathways reliant upon the malate/OAA shuttle within *T. pseudonana* and found no evidence of any such mechanism that could account for changes in CCM enzymes, as had been previously suggested. Continued study of diatom CCMs will serve to not only improve the understanding of how ongoing climate change and ocean acidification will impact these key primary producers, but it will also support emerging interests into their biotechnological applications.

## Supplementary data

The following supplementary data are available at JXB online.

Table S1. Average pH_NBS_ for each treatment replicate.

Table S2. Growth rates and protein extraction efficiencies for all experimental cultures.

Table S3. List of all differentially abundant proteins for the LCHpH and HCLpH comparison.

Table S4. List of all differentially abundant proteins for the LCHpH and LCLpH comparison.

Table S5. Full statistical output for the LCHpH and HCLpH comparison.

Table S6. Full statistical output for the LCHpH and LCLpH comparison.

Supplemental information: Detailed laboratory procedures, experimental apparatus construction, and extended lists of software settings.

## Acknowledgments

The authors thank David Sleat, Haiyan Zheng, and Amenah Soherwardy for their expertise in completing proteomic sample preparation and LC-MS/MS protein sequencing, and for advising on the ensuing data analysis.

## Author contributions

ABK: conceptualization; ABK: funding acquisition; ARH and ABK: methodology; ABK and ARH: investigation; ARH: formal analysis; ARH: data curation; ARH: writing - original draft; ABK and ARH: writing - review & editing; ARH: visualization; ABK: supervision.

## Data availability

The raw mass spectrometry files and DIA-NN main report file have been deposited to the ProteomeXchange Consortium (http://proteomecentral.proteomexchange.org) via the PRIDE partner repository (Perez-Riverol *et al*., 2025) with the dataset identifier PXD064793.

## Abbreviations

AOX: mitochondrial alternative oxidase
CA: carbonic anhydrase
CCM: carbon concentrating mechanism
CET: cyclic electron transport
ETC: electron transport chain
FDR: false discovery rate
HCLpH: high pCO_2_/low pH
LCHpH: low pCO_2_/high pH
LCLpH: low pCO_2_/low pH
MDH: malate dehydrogenase
ME: malic enzyme
NDH: NADH-like complex
OAA: oxaloacetate
OMT: 2-oxoglutarate/malate translocator
PEP: phosphoenolpyruvate
PEPC: phosphoenolpyruvate carboxylase
PEPCK: phosphoenolpyruvate carboxykinase
PGR5: proton gradient regulation 5 protein
PGRL1: PGR5-like photosynthetic phenotype 1 protein
PTOX: plastidial terminal oxidase
PYC: pyruvate carboxylase
SDR: short-chain dehydrogenase/reductase family
SLC4: solute carrier family 4

ALT TEXT: Venn diagrams of the total number of proteins that were either more or less abundant in each of the two treatment comparisons.

ALT TEXT: Graphs showing the statistical results for all 7,803 proteins that passed quality control filtering in each of the two treatment comparisons evaluated in this study.

ALT TEXT: A visual schematic of the proposed carbon concentrating mechanism within a *T. pseudonana* cell informed by statistically significant differential protein abundance data generated in this study.

ALT TEXT: A zoomed in view of the proposed CO_2_ recapture mechanism indicated by the black box insert in Fig. 3. Schematic shows the process through which CO_2_ released in the mitochondrial matrix can be recaptured and fed back into the CCM.

