## Supplemental information for "Proteomic responses under differing pH and pCO_2_ levels in the diatom *Thalassiosira pseudonana* are consistent with a hybrid carbon concentrating mechanism"

*Cleaning and sterilization*

Polycarbonate bottles and tubes were first soaked in 2% Micro-90 cleaning solution (Cole-Parmer, Vernon Hill, IL, USA) for 72 h before rinsing and soaking in 1 M HCl for an additional 72 h. Bottles and tubes were then rinsed with ultrapure water from a Milli-Q Advantage A10 water purification system (MilliporeSigma, Burlington, MA, USA), before being filled with ultrapure water and microwave sterilized (Keller et al., 1988).

*CO2SYS constants*

Within CO2SYS (Lewis and Wallace, 1998) the following constants were used for all calculations: K_1_ and K_2_ dissociation constants for carbonic acid from Lueker et al., 2000, the K_SO4_ dissociation constant from Dickson et al., 1990, the K_HF_ dissociation constant from Perez and Fraga, 1987, and the total boron formulation from Lee et al., 2010.

*Experimental apparatus continued*

Three holes were drilled in each 500 mL experimental culture bottle. The first was in the cap of the bottle and was sized to accommodate the Thermo Orion pH electrode. The other two holes were drilled on opposing sides of the neck of each bottle, which were then each fitted with a 2.4 mm and 3.2 mm barbed hose fitting, respectively. The smaller fitting was used for 0.006 M HCl delivery when pH rose above its set point. The larger fitting was connected to a piece of tubing that ran to the bottom of the bottle and was used to bubble a mixture of ultra zero grade air and CO_2_ (Airgas, Radnor, PA, USA) through the entire culture volume to fully mix each acid addition. The small peristaltic pumps in conjunction with the 2.4 mm barbed hose fittings ensured acid was delivered in small steady drops to allow ample time for cultures to mix without overshooting the target pH. Gas mixtures passed through 0.2 μm sterile cartridge filters before reaching the experimental cultures in order to minimize potential bacterial contamination. Experimental cultures could not be kept sealed due to the aeration, which allowed ambient indoor air to diffuse into the culture headspace over time. The improvement in pH control observed after the implementation of acid-independent aeration every 15 min was, therefore, likely the result of reducing interactions with ambient indoor air, which can be >200 ppm CO_2_ higher than current atmospheric levels (Persily and Gorfain, 2009). The observed variability in pH after this modification was within the manufacturers reported precision for all instrumentation used to control culture pH, and therefore, could not have been further improved upon using the system described here. During testing and experimentation with the revised apparatus, pH was monitored throughout each day and no large pH excursions, like those seen with the original system, were observed for any culture.

*Proteomic sample processing*

Proteomic sample preparation was conducted following the SP3 method (Hughes et al., 2014). For each sample, 10 μg total protein was diluted to a final concentration of 0.1 μg μL^-1^ in buffer containing 50 mM HEPES pH 8.0, 1% SDS, 50 mM EDTA, and 10 mM DTT. Samples were denatured at 60°C for 30 min and briefly centrifuged to remove condensation. Iodoacetamide (IAA) was then added to a final concentration of 40 mM to alkylate samples and left to incubate for 1 h in the dark. A 1:1 mixture of Sera-Mag Carboxylate SpeedBeads E3 and E7 (Cytiva, Marlborough, MA) was added to each sample to a final ratio of 10:1 bead mixture to total protein. Ethanol was then added to a final concentration of 50% and samples were mixed for 5 min at 1200 rpm. Sample tubes were placed in a magnetic stand to collect the now protein coated beads on the side wall of the tube before the supernatant was discarded and the beads were washed with 80% ethanol. Residual ethanol was briefly allowed to evaporate before digestion buffer (100 mM NH_4_HCO_3_, 2 mM CaCl_2_, 1:25 trypsin) was added and samples were left to incubate at 37°C for 1 h while mixing at 1000 rpm. After digestion, sample supernatant was transferred to a bead free microcentrifuge tube and acidified to a pH of 3 with 10% formic acid. Samples were next desalted using C18 stage tips before a 3-phase peptide elution with 15%, 30%, and 50% acetonitrile. Samples were then concentrated under vacuum before peptide sequencing with a Thermo Orbitrap Eclipse Tribrid mass spectrometer with a Dionex U-3000 Rapid Separation nano LC system. Proteomic sample preparation and sequencing were completed by staff at the Biological Mass Spectrometry Facility of Robert Wood Johnson Medical School and Rutgers, The State University of New Jersey.

*In silico spectral library preparation*

A library of known *T. pseudonana* protein sequences had to first be created before an in silico spectral library could be produced. All protein sequences available through UniProt for *T. pseudonana*, which at the time of creation was 11,934 sequences, were used to create the initial reference library. Most sequences available under taxonomic ID 35128 were generated from *T. pseudonana* CCMP 1335, but some were generated from wild populations and other clonal isolates (Alverson et al., 2007; Gastineau et al., 2021). While a unique taxonomic ID exists specifically for CCMP 1335 (296543), it has seen little use, containing only 15 sequences. Therefore, all protein entries that did not clearly originate from CCMP 1335 were removed from the reference library to prevent the potential inclusion of protein variants produced from alleles that are not present within this strain. Upon initial testing with DIA-NN, it became apparent that numerous *T. pseudonana* CCMP 1335 proteins have duplicate entries within UniProt. DIA-NN is capable of recognizing and merging similar entries into one protein group for quantification; however, for *T. pseudonana* this was relatively rare, resulting in the classification of many peptides as nonproteotypic. To prevent the mischaracterization of peptides that could hinder accurate protein quantification, duplicate entries were manually removed. Protein entries based on either transcriptomic evidence or targeted protein sequencing were retained over entries predicted from genomic sequences alone. 11,661 protein entries remained after all filtering was complete. These entries along with sequences from a universal contaminants library (Frankenfield et al., 2022) were then used to generate the in silico spectral library. The following DIA-NN (Demichev et al., 2020; Kistner et al., 2023) settings were used during spectral library generation: FASTA digest for library-free search enabled; deep learning-based spectra, RTs and IMs prediction enabled; protease set to trypsin/P; 1 missed cleavage allowed; 1 variable modification allowed with N-term excision, C carbamidomethylation, and Ox(M) modifications enabled; a peptide length range of 7-30; a precursor charge range of 1-4; a precursor m/z range of 300-1800; and a fragment ion m/z range of 200-1800.

*Data analysis with DIA-NN*

Before protein quantification could proceed with the newly created spectral library, DIA-NN was first run with experimental samples classified as unrelated runs to determine the specific mass accuracy and scan window settings for optimal analysis of this specific dataset. This optimization resulted in an MS1 accuracy of 5.0 ppm, an MS2 accuracy of 14.0 ppm, and a scan window radius of 5. These settings were then fixed for the full analysis run.

For protein quantification, DIA-NN was supplied the original FASTA sequences for both the curated *T. pseudonana* reference library and the universal contaminants library, the in silico spectral library, and all 12 experimental data files. DIA-NN was then run using the high precision Quant UMS quantification strategy with the following settings: a precursor FDR of 1%, MBR enabled, and protein inference enabled. All other parameters were set to the recommended default settings in the DIA-NN user manual.

The DIA-NN main report file was next imported into R ver. 4.4.1 (R Core Team, 2024) for quality control filtering. DIA-NN output was first filtered to remove proteins that did not meet the following criteria: library q value ≤ 0.01, library protein group q value ≤ 0.01, protein group q value ≤ 0.01, and protein group posterior error probability ≤ 0.2. The quality control statistics generated with the high precision Quant UMS method were used to remove low confidence protein quantifications and were set as follows: Quantity.Quality ≥ 0.7, Empirical.Quality ≥ 0.5, and PG.Max.LFQ.Quality ≥ 0.7. Any protein names containing the unique prefix associated with the universal contaminants library were removed. Lastly, proteins that appeared in less than three samples per treatment were removed to ensure only reliable statistical contrasts were interpreted. Filtered data were then passed into MSqRob2 Bioconductor R package ver. 1.12.0 (Goeminne et al., 2016; Goeminne et al., 2020; Sticker et al., 2020) where protein intensities were log_2_ transformed before a robust ridge regression model was fit.
